# Ecosystem engineer restores trophic interactions: beavers alter landscape of risks and resources for megafauna

**DOI:** 10.64898/2026.09.16.751978

**Authors:** Izabela Fedyń, Michał Ciach

**Affiliations:** Department of Forest Biodiversity, Faculty of Forestry, University of Agriculture in Krakow, al. 29 Listopada 46, 31-425 Krakow, Poland

**Keywords:** mammal community, trophic interaction, habitat modification, resource–risk trade-off, spatial ecology, camera trap

## Abstract

Ecosystem engineers hold potential to modify the environment and resource availability. The recovery of the Eurasian beaver reintroduces both a novel trophic component, as herbivore and prey, and an ecological driver that alters landscape of resources and risks. Together, the species’ comeback may trigger cascading effects on the ecosystem and shape relationships among megafauna, with contrasting effects on different trophic groups. We studied 30 beaver sites and paired reference sites in forests across Poland to assess how beaver presence affects habitat use and interactions among ungulates, large carnivores and mesocarnivores. With the same species composition and species richness, the number of detections of animals was lower at beaver sites, whereas detection duration was longer there. Structural equation modelling showed that beaver affects mammal community via a combination of direct and indirect pathways. The species presence had a positive direct effect on large carnivores, partly offsetting the negative indirect pathway mediated through reduced ungulate presence. These effects extended indirectly further to mesocarnivores, detection of which was positively related with increasing number of detections of ungulates and large carnivores. Beaver effects on habitat were spatially structured: canopy openness and deadwood abundance declined with increasing distance to water. Ungulates showed spatially diversified response at beaver sites, with the number of detections increasing away from the water, whereas large carnivores were detected most frequently near the water. Ungulates also changed seasonally their behaviour along water-land gradient, indicating a trade-off between resource acquisition and perceived predation risk. Our results show that beaver engineering reorganizes ecosystem through spatial and temporal changes in habitat use, behaviour and trophic interactions among large and medium-sized mammals. Beaver recovery may therefore restore ecological complexity by generating fine-scale mosaics of resources and risk that propagate across multiple trophic levels.

## Introduction

Historically, megafauna played a central role in structuring ecosystems across most terrestrial biomes via herbivory and predation, often generating strong top-down effects (Czyżewski et al., 2026; Malhi et al., 2016). However, global declines in the abundance and biomass of large mammals have simplified many contemporary ecosystems relative to historical conditions (Svenning et al., 2016). The decline or disappearance of large mammals has been associated with far-reaching ecological consequences, including altered vegetation dynamics, disrupted trophic interactions and the loss of ecosystem processes that were previously maintained by activity of megafauna (Galetti et al., 2018). Such simplification is now recognized as a fundamental dimension of anthropogenic change, with consequences extending beyond biodiversity patterns to ecosystem functioning and stability trough cascading effects across trophic levels (Estes et al., 2011).

In recent decades, however, parts of Europe have experienced a recovery of several large mammal species, including apex predators and large herbivores, whose key ecological roles were previously diminished or lost across much of their former range (Deinet et al., 2013). The recovery of large-bodied species has stimulated growing interest in the ecological consequences of species return and the extent to which restored populations may contribute to trophic rewiliding through the reestablishment of lost biotic interactions (Svenning et al., 2016; Trouwborst & Svenning, 2022). Although trophic rewilding is often discussed in the context of large herbivores and carnivores, recovering medium- or even small-bodied species that hold potential for ecosystem engineering may contribute disproportionally through complementary pathways by reshaping the physical template within which species interactions occur (Byers et al., 2006; Sanders et al., 2014). Nowadays, one of the most rapidly expanding mammals is Eurasian beaver (*Castor fiber*), which recolonize European landscape across the entire gradient of anthropogenic disturbance (Ciach et al., 2023; Halley et al., 2021). The Eurasian beaver is an ecosystem engineer capable of generating extensive environmental modifications (Jones et al., 1994). Through hydrological alterations and vegetation changes beavers profoundly reshape habitat structure and heterogeneity (Kivinen et al., 2020; Law et al., 2017). Beaver-modified sites are recognized as biodiversity hotspots, supporting diverse assemblages of plants, invertebrates, birds and mammals (Law et al., 2016, 2019; Nummi & Holopainen, 2020). Although beaver activity is most often framed within aquatic habitats, consequences of its presence extend into adjacent terrestrial ecosystems and affect terrestrial biota (Fedyń et al., 2024; Nummi et al., 2011).

The recovery of the European beaver reintroduces not only an ecosystem engineer that modifies habitats, but also a species that participates in trophic interactions within mammal communities. As herbivore, beaver consumes woody vegetation and may locally compete with large herbivores for plant food resources (Svanholm Pejstrup et al., 2023). Simultaneously, beaver constitutes a potential prey for large carnivores (Gable et al., 2018), thereby the species comeback to the ecosystem reestablish direct trophic links that had been absent or weakened for centuries. Additionally, beaver-induced changes in habitat structure may indirectly reshape trophic interactions by altering the distribution of food, cover and perceived predation risk, and consequently the probability of encounter between prey and predator. Canopy openings and regenerating vegetation may increase forage availability and attract herbivores (Wilson et al., 2024), whereas dense vegetation and accumulated deadwood may reduce visibility and constrain movement or escape opportunities. Because resource availability and predation risk jointly shape species distribution, habitat selection and behaviour (Hodson et al., 2010; Laundré et al., 2010; Schmidt & Kuijper, 2015), beaver engineering may generate substantial indirect effects on mammal communities. Beaver-modified habitats may form resource–risk mosaic, in which the benefits and costs of habitat use differ among species. Therefore, the final effects of beaver presence on individual trophic groups reflect not only direct relationships with beavers, but also indirect effects mediated by habitat modification and interactions with other trophic groups.

Seasonal changes in beaver activity and its uneven intensity within territories are likely to make spatially and temporally diversified effects on mammal communities. Beaver sites are characterized by pronounced small-scale spatial heterogeneity in vegetation composition and structure, resulting from tree felling, flooding and subsequent succession of vegetation (Kivinen et al., 2020). As central-place foragers associated with fixed territories, beavers concentrate their activity around shorelines, lodges and repeatedly used foraging routes (Fryxell & Doucet, 1991). The intensity and form of habitat modification change with distance from water, producing fine-scale gradients in canopy openness, deadwood accumulation and regeneration of vegetation. Such gradients may determine which parts of beaver-modified sites are used by different mammal groups and where interactions among them occur. Beaver effects also vary temporally, as seasonal changes in the species activity alter both the intensity of habitat modification and the availability of beavers as prey (Sovie et al., 2025). Consequently, the strength and spatial distribution of beaver-mediated effects on terrestrial mammals may shift throughout the year and within small-scale habitat patches.

Despite the ongoing recovery of the Eurasian beaver across much of its former range, its role in shaping the organization and functioning of terrestrial mammal communities remains poorly understood. Evidences from North America demonstrate that interactions involving beavers can scale up beyond pairwise trophic relationships: wolf predation alters beaver foraging behaviour and thereby affects the composition and structure of forests surrounding wetlands (Gable et al., 2023). Therefore, novel trophic interactions and reshaped distributions of resources and predation risk may influence terrestrial mammal communities through both trophic- and habitat-mediated pathways. Because these pathways may operate in opposing directions and differ among trophic groups, their combined consequences for mammal activity, behaviour and spatial distribution remain difficult to predict. The aim of this study is to assess how the presence of the Eurasian beaver influences the community composition, space use and behaviour of megafauna in temperate ecosystem. Specifically, we examine responses of ungulates, large carnivores and mesocarnivores to beaver-modified habitats and evaluate the direct and indirect pathways linking beaver presence, habitat modification and interactions among studied trophic groups. We hypothesized that beaver effect would be group-specific, reflecting differences in direct trophic links with beavers and in responses to beaver-driven changes in habitat. We further expected these effects are spatially and temporally diversified, because both the intensity of beaver activity and the resulting habitat modifications are unevenly distributed within sites and throughout the year. By considering both beaver-induced habitat modification and trophic relationships among mammals, we tested how beaver recovery reorganizes interactions within terrestrial mammal communities.

## Methods

### Study area

The study was conducted in Poland, encompassing lowland, upland and mountainous landscapes typical of the temperate climatic zone of Central Europe. Lowland plains dominate the north and central regions, while uplands and mountain ranges such as the Carpathians and Sudetes stretch along the southern border. Forests cover around 30% of the country’s area, while agricultural land accounts for 60%. Urban areas cover 5% and Poland’s population is about 37.5 million as of 2024, resulting in an average population density of 120 people per km².

Study sites were located in areas representative of the altitudinal gradient – from lowland forests situated ca 200 m above sea level in northeastern Poland, through upland zones ranging from 200 to 400 m, to mountainous regions reaching up to 600 m in elevation. Lowland sites were located within deciduous and coniferous forests dominated by species such as hornbeam (*Carpinus betulus*), pedunculate oak (*Quercus robur*) and Scots pine (*Pinus sylvestris*). Upland areas featured mixed land use, with forests interspersed with agricultural fields, while mountainous locations were primarily covered by stands of European beech (*Fagus sylvatica*) and silver fir (*Abies alba*). Riparian habitats along streams were naturally dominated by alder (*Alnus* spp.), willow (*Salix* spp.) and poplar (*Populus* spp.).

During the study period, mean annual air temperature in Poland was 9.5°C (IMGW-PIB, 2023). Annual precipitation totalled 534.4 mm in 2022 (IMGW-PIB, 2023). Seasonal dynamics was well-marked in the study area, reflecting coordinated changes in temperatures and vegetation phenology. In 2022, December was the coldest month, with a mean air temperature of 0.4°C and snow cover persisted for 75 days in mountainous regions (Lesko) in the southern part of the study area and for 52 days in lowlands (Białystok) in the northern part of the study area (IMGW-PIB, 2023). Spring was characterized by rapidly increasing temperatures and vegetation development started in March. These conditions progressed into summer, when vegetation productivity peaked and thermal conditions were highest, with August reaching a mean temperature of 20.5°C. Autumn was marked by progressive cooling and vegetation senescence, leading into leafless the winter period.

Today, Poland is experiencing the recovery of an almost extinct beaver population and hosts one of the largest and most stable populations of this species in Europe. The population is estimated at over 120000 individuals, widely distributed across diverse habitats ranging from forested wetlands and agricultural landscapes to urban watercourses (Ciach et al., 2023).

### Field methods

#### Study sites

The study was conducted across diverse patches of temperate forests and associated mammalian communities (Fig. 1). The research design includes 30 active Eurasian beaver sites (hereafter: beaver sites) and 30 corresponding reference stream sections without signs of beaver activity (hereafter: reference sites). Beaver sites were defined as stream sections inhabited by beaver family with visible species’ activity, mainly through dam construction and vegetation alteration, with a minimum length of 200 m and at least several years of active beaver presence. Reference sites were either part of the same stream or similar streams of the same order, located within similar habitat and landscape. Each reference site was situated at least 1 km away to prevent potential overlap due to beaver expansion during the study period. Selected pairs of sites were separated by a minimum of 5 km to increase independence of studied mammal communities and include different landscape contexts.

**Fig. 1.**
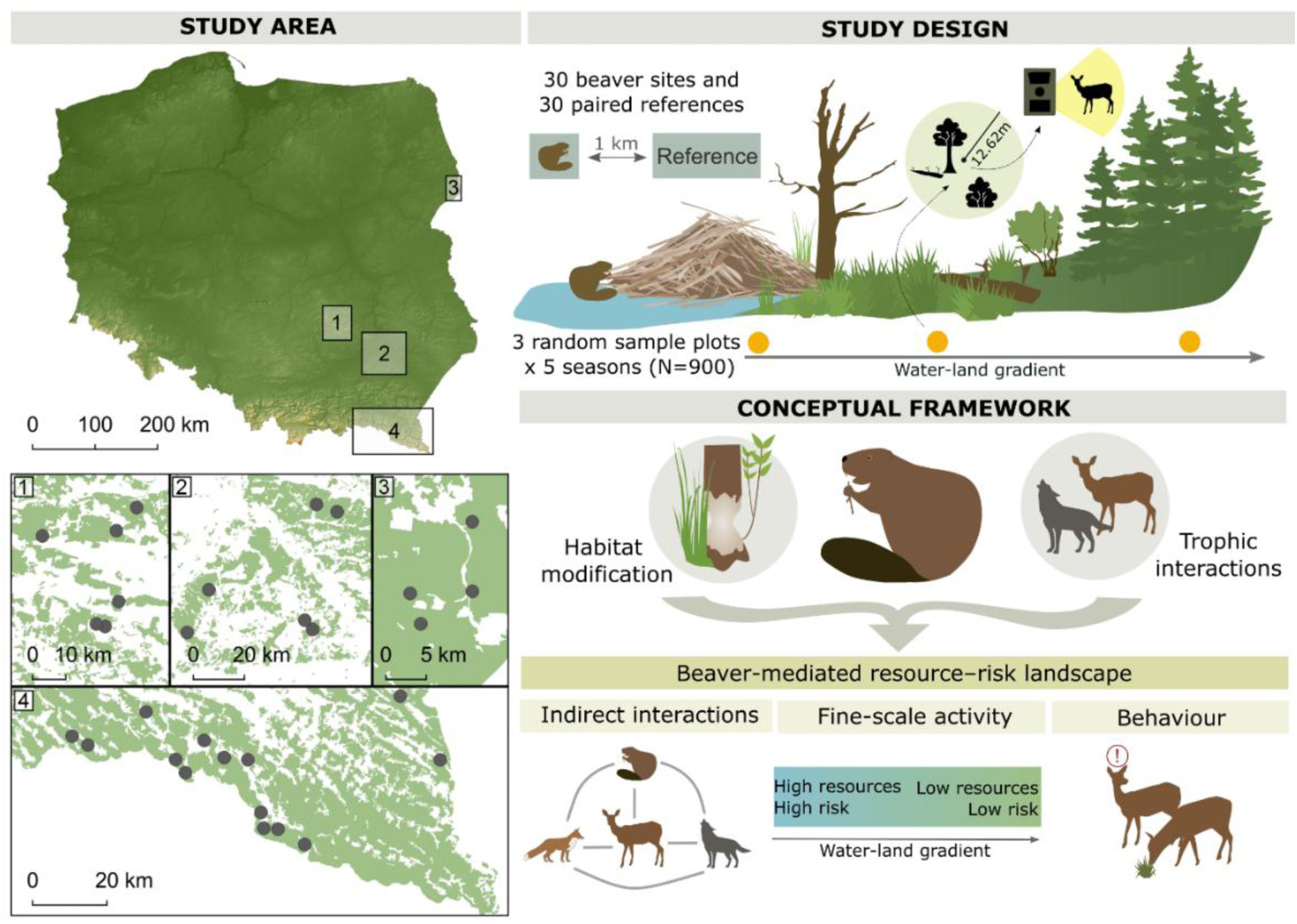
The location of four study regions in Poland (Central Europe), including the locations of the 30 paired study plots (grey dots), each consisting of a Eurasian beaver (*Castor fiber*) site and a paired reference site. The study design illustrates the study design and conceptual framework summarizes the hypothesized pathways linking beaver-mediated habitat modification with trophic interactions through changes in habitat use and animal behaviour.

#### Selection of random sampling plots and camera deployment

Data collection was carried out between September 2022 and August 2023 to cover seasonal changes in mammal activity and distribution (Kays et al., 2020). Five sampling periods (hereafter: seasons) were selected to represent contrasting phenological phases of the annual cycle, characterized by seasonal changes in vegetation development, food availability, climatic conditions. This pronounced seasonal cycle provided the environmental context for temporal variation in mammal activity (Kays et al., 2020). Selected seasons included: early autumn (1.09.2022 – 8.11.2022), autumn (10.11.2022 – 12.12.2022), winter (1.01.2023 – 15.03.2023), spring (1.04 – 15.05.2023) and summer (20.06.2023 – 5.08.2023). Each study site was subdivided into three distance zones from the waterline: (I) < 3 m – shoreline zone; (II) 3–30 m – terrestrial beaver activity zone with feeding trails or felled trees; (III) 30–100 m – area usually beyond beaver activity range (Stoffyn-Egli & Willison, 2011) and without beaver signs. One random location per distance zone was selected at each site in each season, resulting in total 900 sampling plots, where a camera trap was deployed and habitat characteristic was described. During each of season, cameras (Browning Spec Ops HP5) were installed at the sampling plots for approximately two weeks (mean = 16.4 days, median = 14 days, range 7 - 30, IQR 13-17). Deployment duration varied among plots due to logistical constraints, equipment availability and field accessibility, particularly during winter. One camera from reference site was lost resulting in reduced number of sampling plots (N = 899). Variation in sampling effort was accounted in subsequent analyses by including number of camera-days as an offset in the models. Camera traps with detection range of ca. 30 m recorded pictures both day and night, triggered by animal movement with a 5-second delay between consecutive activations. Cameras were mounted on trees at a height of 0.8 to 2 m, adjusted to terrain slope, with the lower picture frame edge aimed at a point 2 m from the lens to standardize detection range. Selected camera placement and direction had to ensure possible broad visibility, avoiding dense vegetation or steep slopes, potentially influencing animals’ movement.

#### Habitat characteristic

At each sampling plot, habitat characteristics were described within a circular plot of 0.05 ha (12.62 m radius) centred on the camera trap location. Measurements of vegetation were collected to assess habitat features potentially influencing mammal occurrence and activity. Canopy openness was visually estimated as the percentage of sky visible through the tree crowns, recorded to the nearest 5%, with higher values indicating a more open canopy. The coverage of understory was assessed by estimating the percentage cover of shrubs and trees with a diameter at breast height (DBH) below 7 cm and height between 1 and 5 m. Ground vegetation cover was estimated by categorizing and recording the percentage contribution (summing to 100%) of grasses and sedges, deciduous saplings up to 1 m high, conifer saplings up to 1 m, herbaceous plants, blackberries (*Rubus* spp.), blueberries (*Vaccinium* spp.) and bare ground or leaf litter. Deadwood was quantified by counting standing dead trees (snags), uprooted trees and fallen dead trees (logs) with DBH at least 7 cm with a minimum height/length of 1.5 m.

### Data handling and analyses

#### Camera trap data processing

All images from the camera traps were first manually screened to remove pictures with no animal recorded. Subsequently, the remaining images were described using Timelapse v.2.3.0.8 software. For each detection, the species, the number of individuals and their behaviour were documented. In the case of photos containing more than one individual, the behaviour of the animal closest to the camera was indicated. Behavioural classification was based on body posture, orientation and visible activity recorded by camera traps. Behaviour was classified into six categories: movement – when the animal was walking or running; vigilance – when body posture indicated alertness or screening; foraging – when the animal was feeding or drinking; rest – when the animal was lying down or engaged in grooming; social interaction – during territorial marking or direct interactions such as mating or playing; unidentified – when the animal was not fully visible and its behaviour could not be precisely determined (description in ethogram in Table S1). In cases where species identification was impossible due to poor image quality or vegetation obstructing key features, the detection was categorized as undefined and excluded from further analyses.

#### Data analysis

Because the camera-trap setup was primarily designed to detect medium- and large-sized mammals, further analyses were restricted to wild mammals belonging to three focal trophic groups: ungulates, large carnivores and mesocarnivores. Of the total dataset comprising 32816 photographs of 25 mammal species, birds, vehicles and humans (Table S2), subsequent analyses included records of terrestrial mammals belonging to distinguished groups, including ungulates: roe deer (*Capreolus capreolus*), red deer (*Cervus elaphus*), fallow deer (*Dama dama*), moose (*Alces alces*), wild boar (*Sus scrofa*) and European bison (*Bison bonasus*); large carnivores: grey wolf (*Canis lupus*), brown bear (*Ursus arctos*) and Eurasian lynx (*Lynx lynx*); mesocarnivores: red fox (*Vulpes vulpes*), European wildcat (*Felis silvestris*), European badger (*Meles meles*), pine marten (*Martes martes*) and raccoon dog (*Nyctereutes procyonoides*).

For each species, the number of recorded pictures was counted and independent detections (hereafter: detections), defined as using a 30-min temporal independence rule: consecutive pictures of the same species at the same camera were assigned to the same detection unless the time gap between photos exceeded 30 min. Taking into account the differences in camera effort, the number of detections was also calculated as the number of the detections of given species on the camera divided by the number of the camera-days and multiplied by 100, resulting in standardized metric of detections per 100 camera-days.

To compare mammal assemblage characteristics and patterns of site use between beaver sites and reference sites, we calculated species richness, number of detections per 100 camera-days and mean detections length for the entire mammal assemblage and separately for the ungulates, large carnivores and mesocarnivores. These metrics were calculated at the site level by pooling data from the 15 camera-trap deployments (sampling plots) conducted across three distance zones and five seasons. Species richness was calculated as the total number of species recorded at least once on any camera within a sampling plot. Number of detections was calculated as the total number of independent detections summed across cameras and seasons and standardized by the total sampling effort, resulting in an activity index expressed as detections per 100 camera-days. To quantify the temporal dimension of site use, for each detection for each species we calculated the number of pictures as a detection length. Mean detection length was then calculated by averaging detection lengths across all detections of species belonging to a given trophic group and across all mammal species for the entire mammal assemblage. Differences between beaver and reference sites in species richness, detections per 100 camera-days and mean detection length were tested for the entire mammal assemblage and for each trophic group using Wilcoxon rank-sum tests.

Differences in mammal community composition between beaver sites and reference sites were assessed using non-metric multidimensional scaling (NMDS) based on Bray–Curtis dissimilarities. The analysis was conducted at the site level (30 beaver sites and 30 reference sites) using a community matrix constructed from the total number of independent detections recorded for each species at each site across the entire study period. The effect of site type on community composition was tested using permutational multivariate analysis of variance (PERMANOVA), and homogeneity of multivariate dispersion between groups was evaluated to verify model assumptions.

For each sampling plot we quantified habitat characteristic, including canopy openness, coverage of undergrowth, the number of snags, logs and uproots, the percentage cover of grasses, herbs, blueberry, blackberry and forest floor diversity expressed using the Shannon index. Differences in individual habitat variables between beaver sites and reference sites were tested using t-tests (Table S3). To reduce model complexity and avoid including multiple partially redundant descriptors of habitat structure, we explored the multivariate structure of vegetation and structural variables using principal component analysis (PCA). Canopy openness showed a strong association with the main ordination axis and represented the transition from closed-canopy forest to more open, beaver-modified patches characterised by greater development of ground vegetation and deadwood presence (Figure S1). To reduce model complexity and avoid including multiple partially redundant descriptors of habitat structure, canopy openness was therefore retained as a single, ecologically interpretable proxy for the local habitat gradient in all subsequent models.

To assess direct and indirect determinants of fine-scale patch utilization disentangle relationships linking beaver activity, habitat characteristic (represented by canopy openness), ungulates, large carnivores and mesocarnivores at the sampling plot scale (N = 900), we applied a piecewise structural equation modelling (SEM; Lefcheck, 2016). The SEM was constructed using a set of a priori defined, ecologically motivated pathways reflecting hypothesised ecological relationships (Table 1). Number of detections were standardized to 100 camera-days and log(x + 1)-transformed, and all continuous variables were subsequently standardized (mean = 0, SD = 1) to facilitate comparison of path-effect sizes across SEM components. All component relationships were then fitted using linear mixed-effects models (LMMs). Beaver presence was included as a binary predictor (1 on beaver sites and 0 on reference sites). All models included random intercepts for sampling units (site × season) to account for hierarchical data structure and potential spatio-temporal non-independence between sampling plots within study site. Following model selection, component models were combined into a single SEM. Global model fit was assessed using Fisher’s C statistic and tests of d-separation. Missing paths indicated by d-separation tests were added when supported ecologically and statistically, resulting in the final model structure. Accordingly, direct paths from ungulates to mesocarnivore activity were included in the final model because the d-separation test indicated a residual dependency between these variables. For each component model, conditional coefficients of determination (R²) were calculated to quantify the variance explained by both fixed effects and random effects. Direct effects were represented by individual path coefficients, whereas potential indirect effects were inferred from sequences of significant directed paths linking variables through one or more intermediate variables.

**Table 1.**
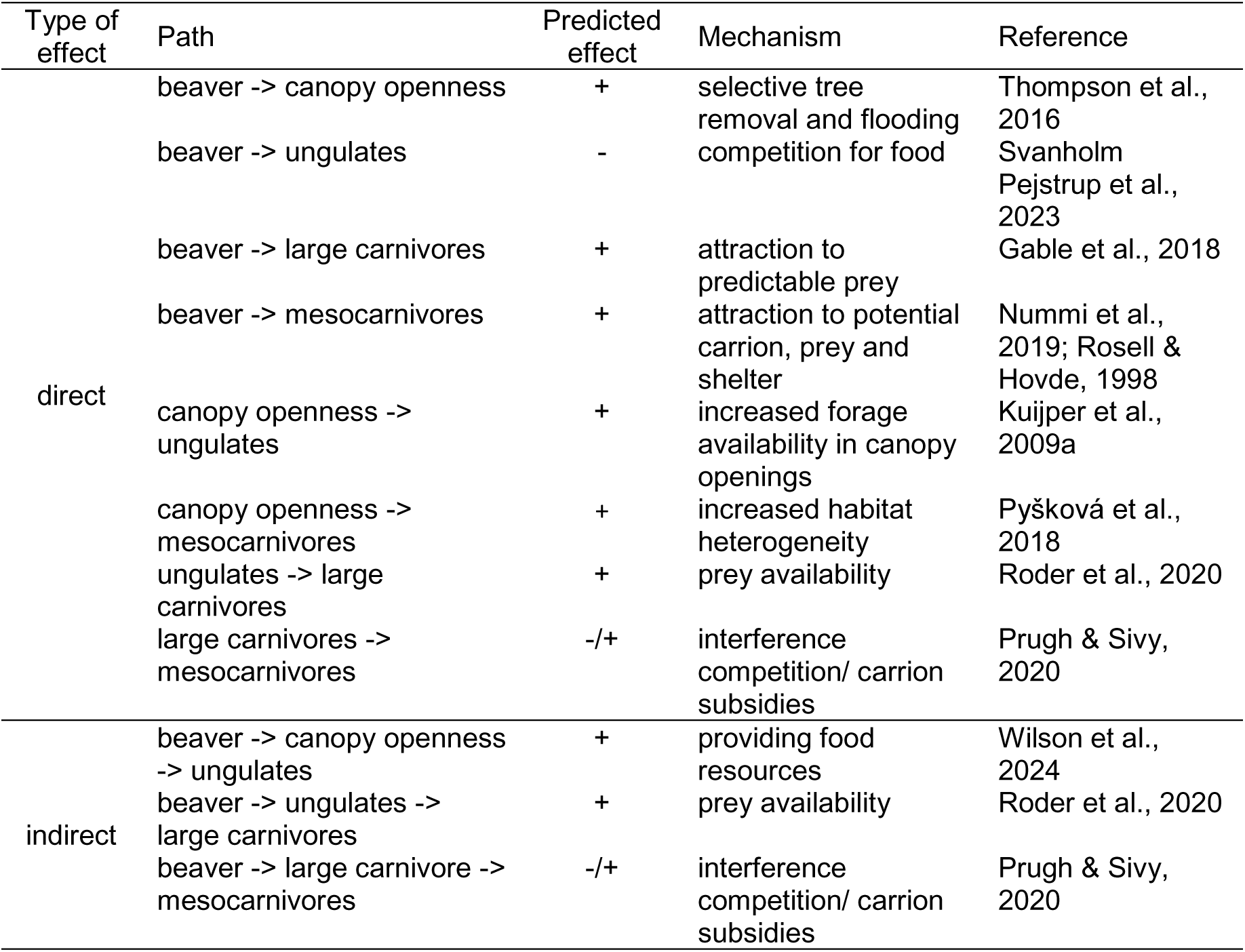
Hypothesized ecological paths included in structural equation model represent pathways through which beaver may influence ungulates, large carnivores and mesocarnivores by altering habitat structure, resource availability and predation risk.

To assess how habitat characteristics and the fine-scale space use of ungulates, large carnivores and mesocarnivores along water-land gradient, we modelled responses as a function of distance to water, site type and their interactions. Canopy openness and deadwood availability were analysed using LMMs, with study site included as a random intercept. For each group, we fitted a generalized LMM with a negative binomial error distribution, where the response variable was the number of detections of the group. Models included distance to water, site type (beaver vs reference) and their interaction, and camera effort (number of camera-days). Variation in sampling effort was accounted for by including the logarithm of camera-days as an offset, and sampling unit (site × season) was included as a random intercept. Model-based marginal predictions were generated along the observed distance-to-water gradient separately for beaver and reference sites. For graphical presentation, predicted mammal detections were standardized to 100 camera-days.

Because animal presence alone may not fully capture response to beaver-induced habitat modifications, we additionally examined behavioural responses of ungulates. For foraging, vigilance and movement, we quantified their proportional contribution to the time budget within each season as the proportion of photographs assigned to each behaviour among all photographs classified into these three categories. We modelled the probability that an observed behaviour was classified as vigilance, foraging or movement using generalized LMMs with a binomial error distribution, with each behavioural category was modelled against all other behaviours. Explanatory variables included site type (beaver vs. reference), distance to water, canopy openness and the number of large carnivore detections at each sampling plot. An interaction between site type and distance to water was included to test whether behavioural responses along water-land gradient differed between beaver and reference sites. Models were fitted both to the pooled dataset for general patterns and separately for individual seasons to assess seasonal variation in behavioural responses. Predicted probabilities and confidence intervals were derived from the fitted models and used to visualize behavioural responses along the water-land gradient. Sampling plot was included as a random effect to account for repeated observations within sites.

## Results

### Community characteristic

A total of 26356 photographs were recorded, representing 6829 independent 30-min detections, with a total sampling effort of 14708 camera-days across 30 beaver sites and 30 reference sites. In total, 14 mammal species were detected, including six ungulates, three large carnivores and five mesocarnivores (Table S2). All species were recorded at both beaver sites and reference sites (Table S2) and the mean number of species did not differ between beaver sites and reference sites for the entire assemblage or for ungulates, large carnivores and mesocarnivores (Fig. 2). Mammal activity, expressed by the mean number of detections was higher at reference sites. In contrast, mean detection length was greater at beaver sites, both for entire assemblage and separately for ungulates, large carnivores and mesocarnivores (Fig. 2). NMDS ordination showed overlap in mammal community composition between beaver and reference sites. PERMANOVA indicated no effect of site type on community composition (F = 0.37, R² = 0.01, p = 0.916) and multivariate dispersion did not differ between groups (p = 0.177).

**Fig. 2.**
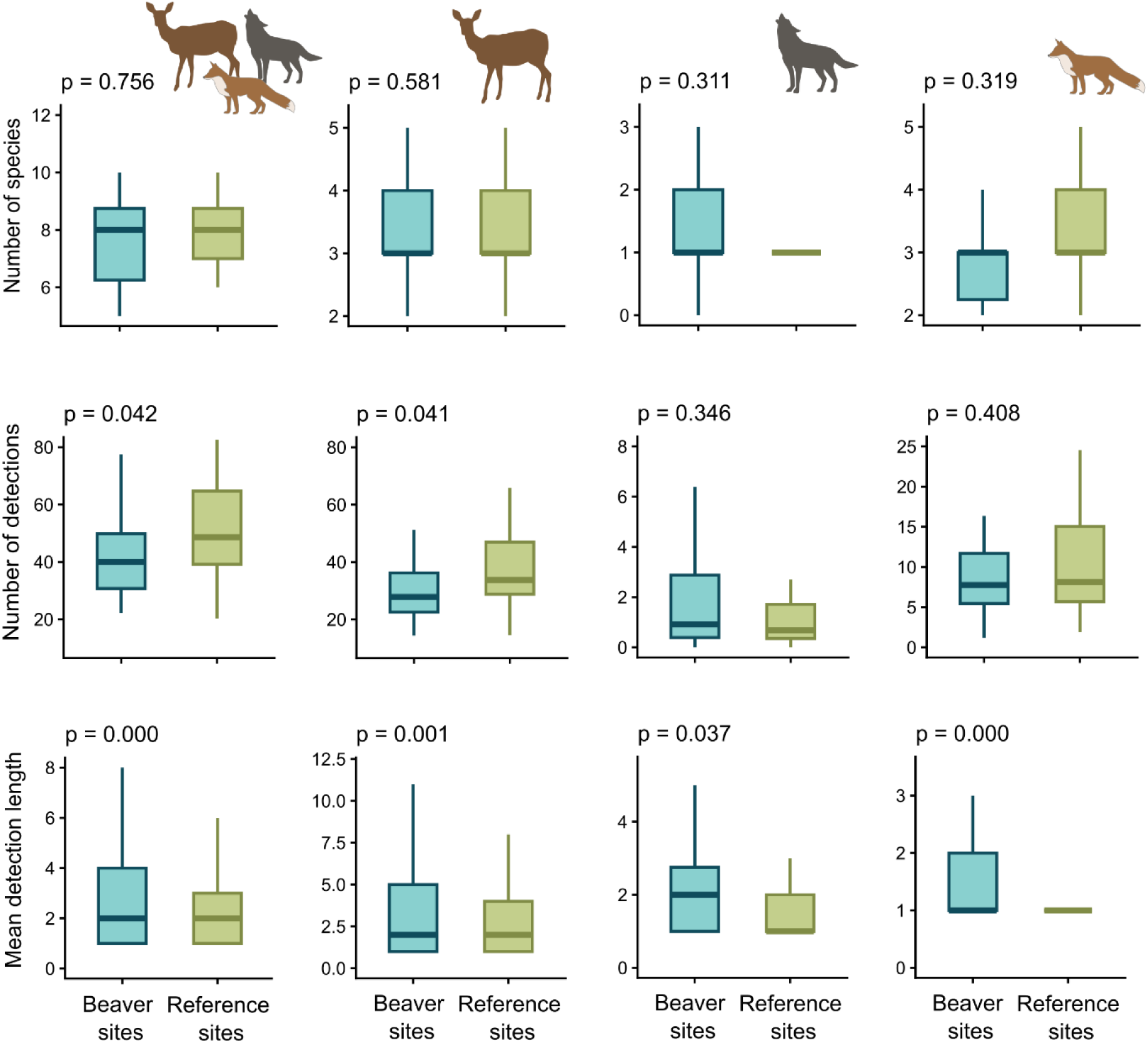
Differences in number of species, number of detections and detection length of mammals on Eurasian beaver (*Castor fiber*) (N = 30) and reference site (N = 30) in Poland (central Europe). Medians, interquartile ranges (IQR) and whiskers extended to 1.5 × IQR are shown. Differences tested with Wilcoxon tests.

### Habitat characteristic

Several habitat characteristics differed between beaver and reference sites (Table S2). Beaver sites had a significantly more open canopy, higher numbers of snags and logs, greater forest-floor diversity and higher cover of grasses and blueberry, whereas herbaceous plant cover was lower. No differences were found in undergrowth cover, the number of uprooted trees, or blackberry cover. PCA revealed a common habitat gradient in which increasing canopy openness was associated with greater deadwood accumulation and corresponding changes in ground-layer vegetation (Fig. S1).

### Cascading effects

The SEM examining relationships among beaver presence, canopy openness, and the number of detection of ungulates, large carnivores and mesocarnivores showed good global fit to the data (Fisher’s C = 4.01, df = 2, p = 0.135). The component models explained 35% of the variation (conditional R²) in canopy openness, 28% in ungulate activity, 30% in large carnivore activity and 13% in mesocarnivore activity (Fig. 3). Beaver presence increased canopy openness (β = 0.53, p < 0.001) and had a negative direct effect on the number of ungulate detections (β = −0.24, p = 0.005; Fig. 3). Canopy openness was also negatively associated with number of ungulate detections (β = −0.09, p = 0.014; Fig. 3), resulting in an additional negative indirect pathway from beaver presence to ungulates. Ungulate detections were positively associated with large-carnivore detections (β = 0.09, p = 0.005; Fig. 3). Beaver presence also had a positive direct effect on large carnivores (β = 0.17, p = 0.043; Fig. 3), but this was partly offset by negative indirect effects mediated through reduced ungulate detections. Mesocarnivore detections were positively associated with both ungulate (β = 0.12, p < 0.001) and large-carnivore detections (β = 0.14, p < 0.001; Fig. 3).

**Fig. 3.**
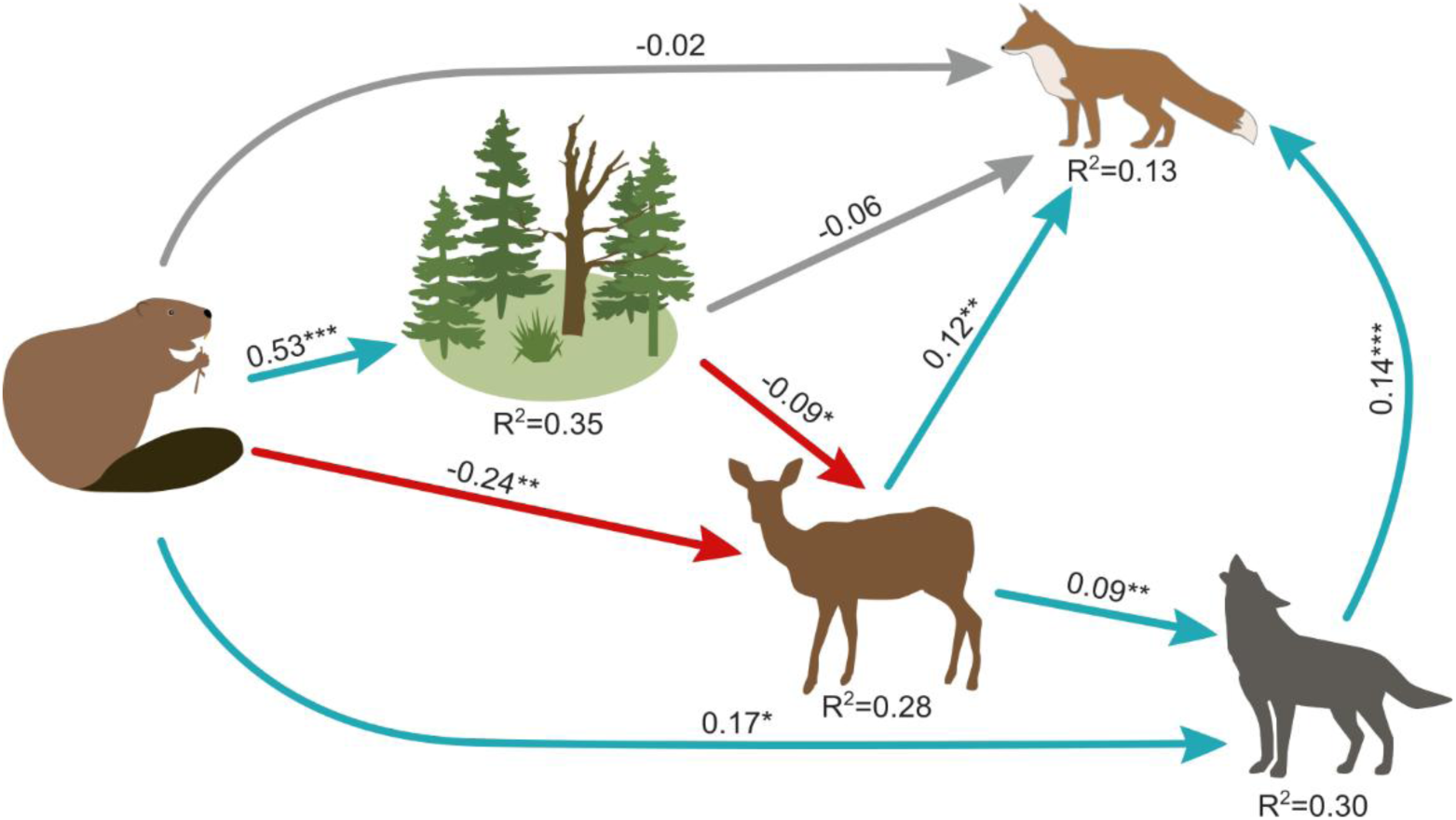
Structural equation model describing the pathways linking beaver presence, canopy openness and the number of detections of ungulates, large carnivores and mesocarnivores on sampling plots on Eurasian beaver (*Castor fiber*) sites (N = 450) and paired reference sites (N = 449) in Poland (central Europe). The model structure was based on the a priori hypothesized pathways (see Table 1). Arrows indicate directional relationships between variables and values next to arrows represent standardized path coefficients. Blue, red and grey arrow indicates positive, negative and non-significant relationship, respectively. Conditional R² values indicate the proportion of variance explained by each component model by fixed and random effects.

### Spatial extent of beaver effects along the water–land gradient

Predicted marginal responses along the water-land gradient showed contrasting patterns between beaver and reference sites (Fig. 4). Canopy openness was highest close to water at beaver sites and declined strongly with increasing distance from the water, whereas no spatial pattern was observed at reference sites. Deadwood abundance also declined with increasing distance from water at both site types, but the decline was substantially stronger at beaver sites (Fig. 4A-B; Table S4). Consequently, differences in canopy openness and deadwood abundance between beaver and reference sites were most pronounced near the water and largely converged at around 70-80 m from the water (Fig. 4A-B). Spatial responses of mammal groups differed along the water-land gradient (Fig. 4C-E; Table S5). Large carnivores detections near the water were more common at beaver sites, but no clear spatial gradient was supported by the models (Fig. 4D; Table S5). In contrast, number of ungulate detections increased with increasing distance from water, with a stronger increase observed at beaver sites (Fig. 4C; Table S5). Mesocarnivore detections showed contrasting patterns between site types, declining with increasing distance from water at reference sites while remaining relatively stable at beaver sites (Fig. 4E; Table S5).

**Fig. 4.**
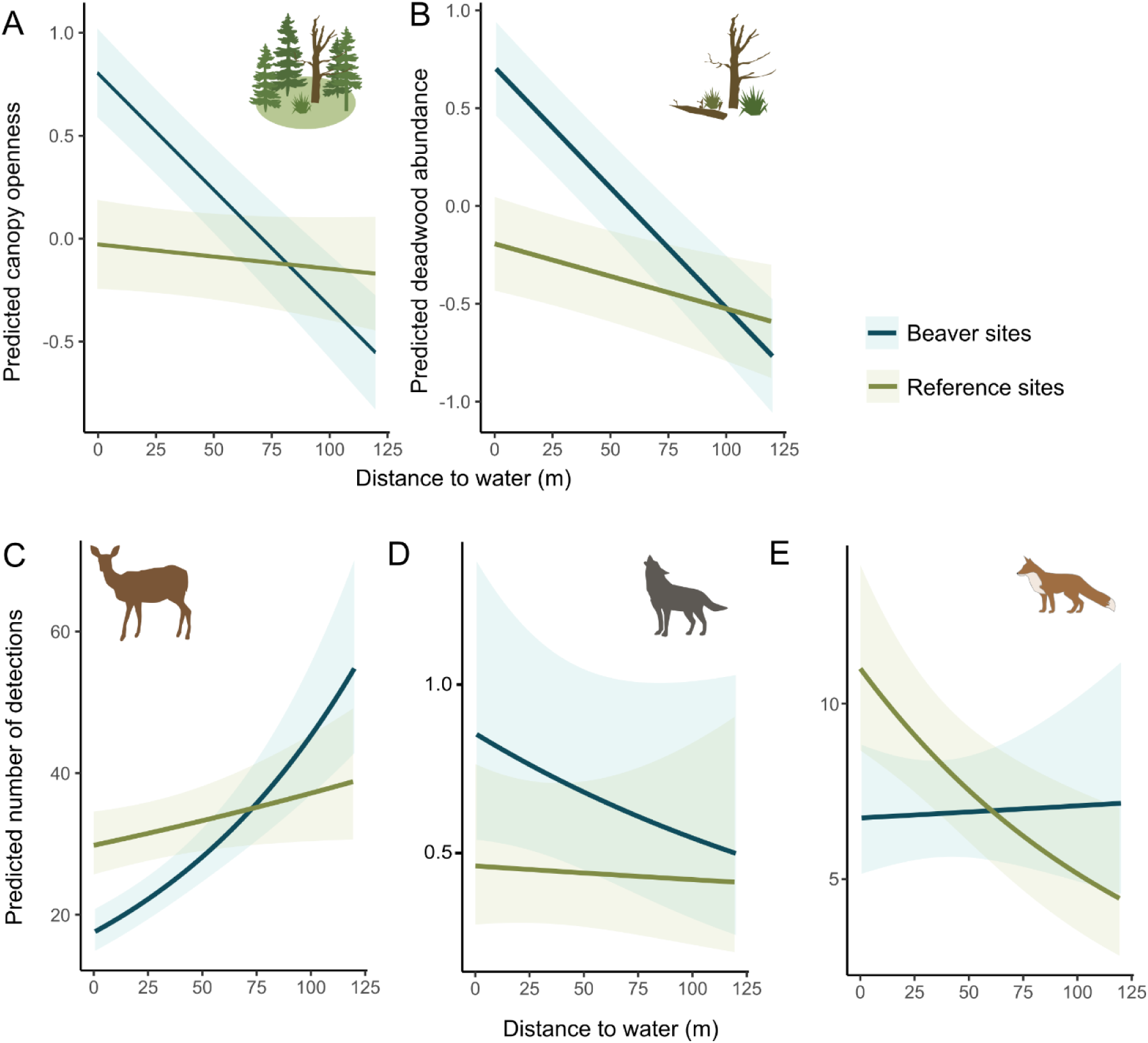
Predicted responses of canopy openness (A), deadwood abundance (B) and number of detections per 100 camera-days of ungulates (C), large carnivores (D) and mesocarnivores (E) along the water-land gradient on sampling plots on Eurasian beaver (*Castor fiber*) sites (N = 450) and paired reference sites (N = 449) in Poland (central Europe). Canopy openness and deadwood abundance are shown on a standardized scale. Lines represent model-based marginal predictions from GLMMs (Table S4 and S5). Shaded areas represent 95% confidence intervals.

Ungulate behaviour varied between beaver and reference sites and changed along the water-land gradient (Fig. 5). At beaver sites, the probabilities of vigilance and foraging were highest close to water and declined with increasing distance, whereas both these behaviours showed the opposite patterns at reference sites (Fig. 5). Movement behaviour showed a contrasting pattern, increasing in greater distances from water at beaver sites but decreasing in greater distances from water at reference sites (Fig. 5). Behaviour was also associated with habitat characteristic and large carnivore occurrence: greater canopy openness was associated with lower probabilities of vigilance and movement, but a higher probability of foraging, whereas a greater number of large carnivore detections was associated with reduced foraging (Fig. 5).

**Fig. 5.**
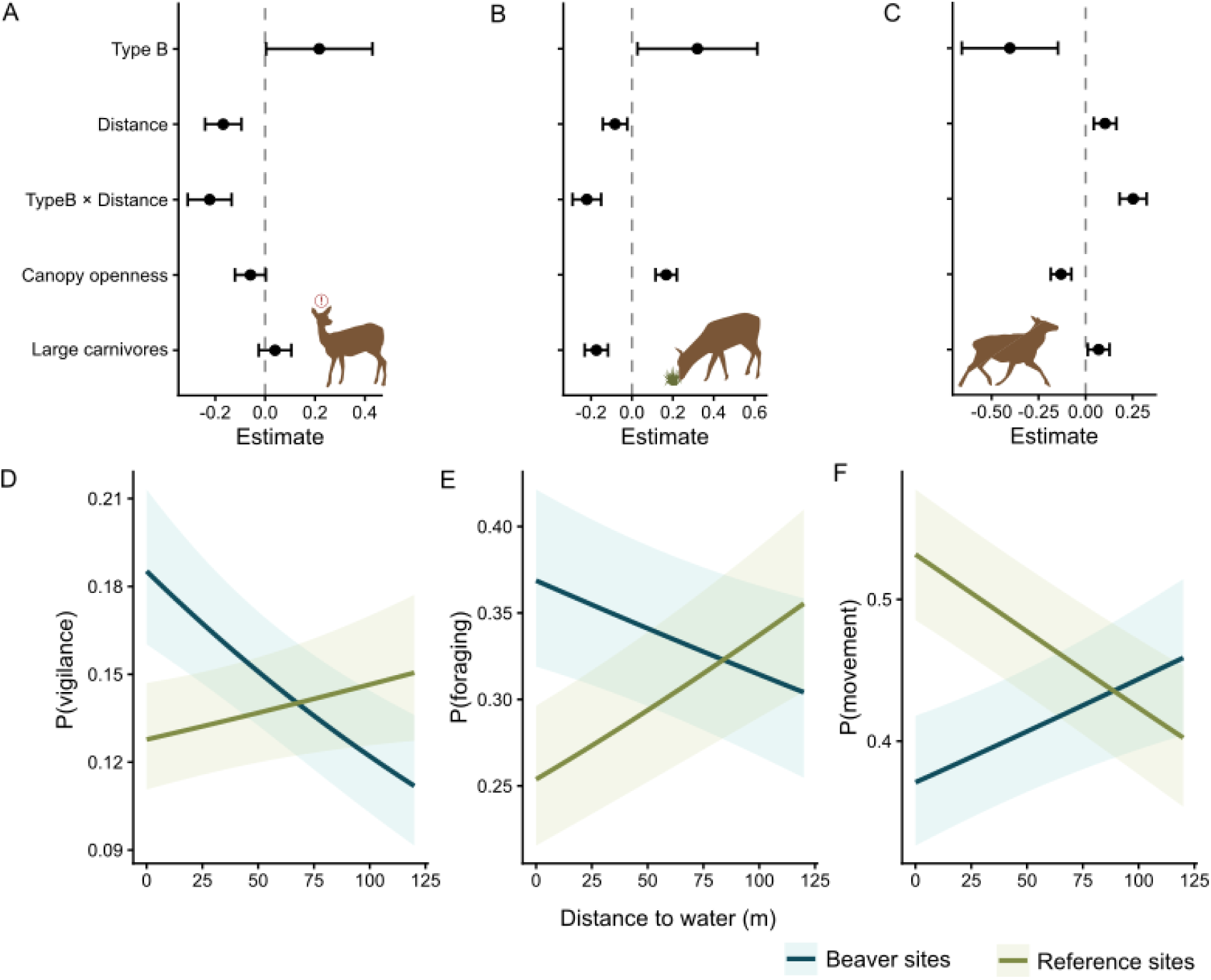
Effects of habitat characteristics on ungulate behaviour. Panels A–C show model estimates (±95% CI) for predictors included in behavioural models for vigilance (A), foraging (B), and movement (C) of ungulates along the distance-to-water gradient on sampling plots on Eurasian beaver (*Castor fiber*) sites (N = 450) and paired reference sites (N = 449) in Poland (central Europe). Model-based marginal predictions (P) from GLMMs are presented for vigilance (D), foraging (E), and movement (F). Shaded areas represent 95% confidence intervals.

The proportional composition of foraging, movement and vigilance remained broadly consistent across seasons, with foraging accounting for 42-53% of records, movement for 33-40%, and vigilance for 14-20% (Fig. 6). Seasonal models revealed marked temporal variation in the relationships of ungulate behaviour with beaver sites, distance from water, canopy openness and large carnivore detections (Fig. 6). During summer, early autumn and autumn, a greater number of large carnivore detections was consistently associated with increased vigilance and reduced foraging (Fig. 6). In summer and autumn, foraging also showed a clear spatial pattern at beaver sites, being concentrated closer to the water and declining with increasing distance from the shoreline (Fig. 6). In early autumn greater canopy openness associated with increased foraging and reduced vigilance (Fig. 6). In winter and spring, greater large carnivores detections were associated with lower vigilance and higher foraging (Fig. 6). In both seasons, vigilance at beaver sites also varied along the water–land gradient, being higher closer to water and declining with increasing distance (Fig. 6). In winter, greater canopy openness was associated with increased vigilance and reduced foraging, whereas in spring it was associated with lower vigilance (Fig. 6).

**Fig. 6.**
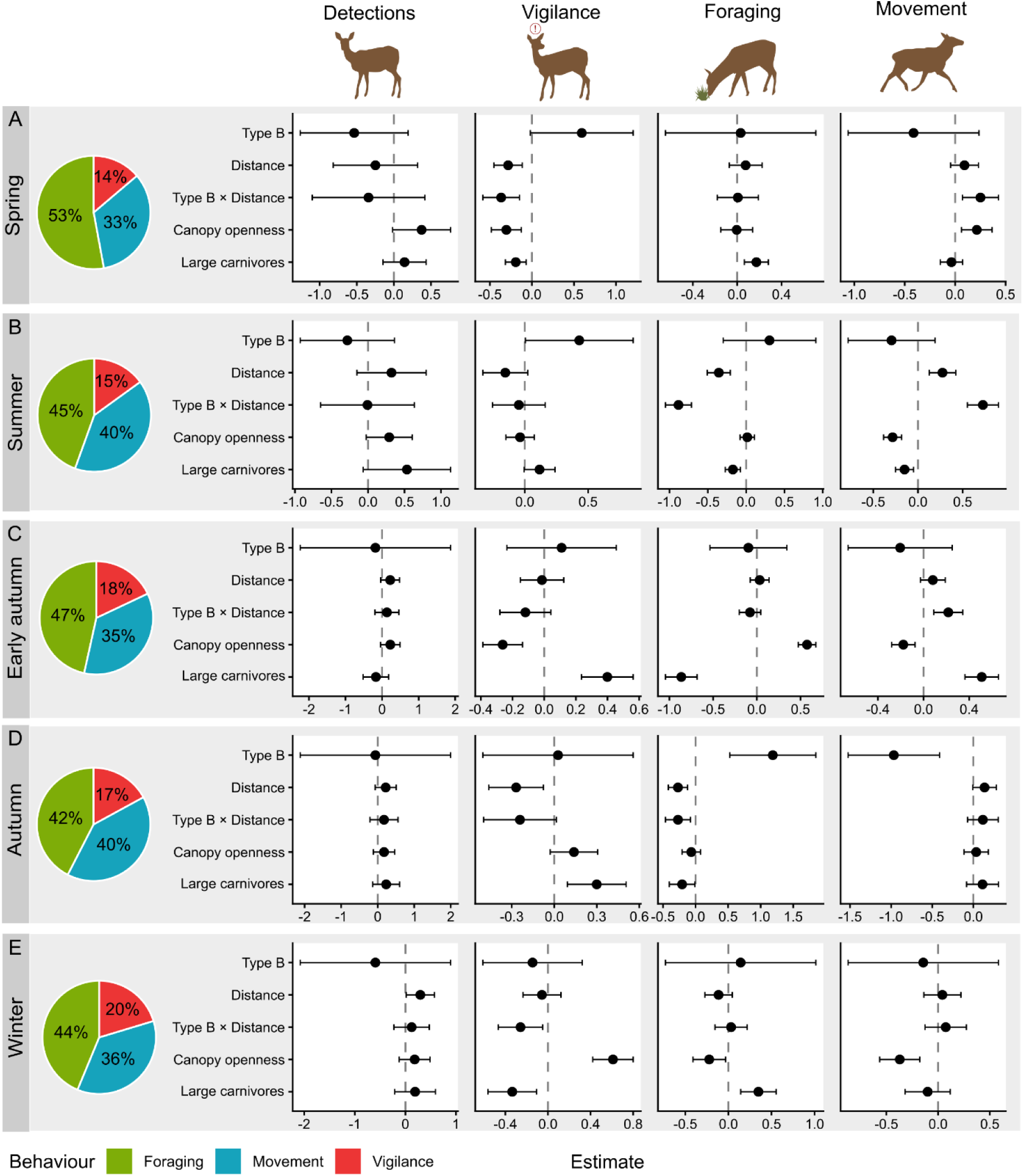
Seasonal variation in ungulate activity and behaviour at Eurasian beaver (*Castor fiber*) sites (N = 450) and paired reference sites (N = 449) in Poland (central Europe). Pie charts show the seasonal composition of behavioural records classified as foraging, movement or vigilance. Coefficient estimates with 95% confidence intervals from GLMMs are presented for the number of detections, vigilance, foraging and movement across spring (A), summer (B), early autumn (C), autumn (D) and winter (E).

## Discussion

Our study shows that beaver presence reorganize the functioning of mammal communities by induced changes in fine-scale space use, animal behaviour and trophic interactions among ungulates, large carnivores and mesocarnivores. These responses apparently emerged from the combined effects of trophic relations with beavers and indirect pathways mediated by habitat modification and the activity of other trophic groups. Together, the ecological consequences of beaver recovery emerge from multiple interacting processes whose relative importance differed among trophic groups and across space and time.

We provide community-level assessment of beaver recovery based on a year-round survey of medium- and large-bodied mammals across multiple temperate forests. Although studied habitats support diverse mammal communities, including several species that remain rare or spatially restricted in Europe, such as wolf, lynx, brown bear, wildcat or European bison, we found no evidence that beaver presence altered species richness or community composition. Previous studies have reported positive effects of beavers on the diversity and activity of terrestrial mammals, particularly in boreal ecosystem, where the high productivity of beaver-modified habitats has been associated with increased use by moose and several mesocarnivores, including red fox, martens and raccoon dog (Nummi et al., 2019). Snow-tracking survey in European temperate forests recorded higher mammal species richness at beaver sites, but this pattern was driven primarily by large carnivores and small mustelids, whereas ungulates and medium-sized carnivores showed no increase in occurrence (Fedyń et al., 2022). Differences among studies may therefore reflect biome, season, taxonomic coverage and the spatial scale at which mammal responses were assessed. As beaver recovery was not expressed by the entrance or loss of species, our results indicate that the ecological importance of the species recovery may emerge through changes in space use, behaviour and relationships among existing trophic groups.

Associations with beaver presence differed markedly among mammal groups. Large carnivore activity was higher at beaver sites, which is consistent with the role of beavers as a predictable prey associated with fixed territories (Gable et al., 2018, 2021). In contrast, ungulate activity was reduced at beaver sites potentially because of overlap in food niche, as both ungulates and beaver rely on woody vegetation (Svanholm Pejstrup et al., 2023). Mesocarnivores showed no correlation with beaver presence, though beaver-related resources may be used by individual species, for example lodges as shelter or individuals as occasional prey (Rosell & Hovde, 1998). However, mesocarnivores were positively associated with large carnivores and ungulates, indicating that direct interactions with beavers are rather opportunistic, but the relation to beaver sites was linked to broader ecosystem context. As facultative scavengers, mesocarnivores can benefit from ungulate carrion and from carcasses generated by large carnivores, which may promote spatial co-occurrence among these groups of predators (Prugh & Sivy, 2020). While ungulates and large carnivores appear to be linked directly to beavers, the response of mesocarnivores demonstrates that beaver recovery can affect trophic groups lacking a strong direct relationship with beavers by modifying interactions among other predators and their prey.

Beaver engineering generated a pronounced fine-scale spatial water-land gradient in both habitat structure and mammal space use that was not observed at reference sites, indicating that these patterns were not induced by water proximity alone. Beavers are central-place foragers whose terrestrial activity is concentrated around lodges, feeding sites and movement routes within approximately 50 m from the shoreline (Graf et al., 2016). This spatial concentration of beaver activity was clearly reflected in pronounced gradients in habitat characteristics, including greater canopy openness and deadwood accumulation near water on beaver sites. Mammal trophic groups responded differently along the same gradient: large-carnivore activity was concentrated near water and declined with distance, whereas ungulate activity increased towards the outer parts of beaver-modified habitat patches. The concentration of large carnivores near water is consistent with their hunting strategies that exploit spatially predictable prey activity, as wolves hunt beavers close to shorelines and near repeatedly used trails and foraging areas, where encounters with beavers are most likely (Gable et al., 2021). Beaver sites may therefore constitute hunting locations within the landscape, which carnivores can exploit by using spatial memory to return to previously learned foraging sites (Gurarie et al., 2022). Additionally, beaver-induced habitat modifications can improve condition of hunting grounds at beaver sites. Accumulation of downed deadwood and reduced visibility due to lush vegetation close to water, may create conditions that restrict movement, escape opportunities or effective vigilance, potentially increasing prey vulnerability to predation. Large carnivores use specific landscape features that constrain prey escape to improve hunting success, including steep terrain and linear barriers (Bojarska et al., 2017; Torretta et al., 2017). From the ungulate perspective, habitat characteristics and higher activity of large carnivores decrease the probability of successful detection of approaching predators, causing these areas to be perceived as risky and imposing additional behavioural costs on ungulates (Halofsky & Ripple, 2008). The combined effects of habitat structure and elevated predator activity near water may therefore explain the reduced use of shoreline areas by ungulates at beaver sites.

Our behavioural models indicates that beaver sites constitute areas of elevated risk, while at the same time providing attractive foraging opportunities, requiring ungulates to balance resource acquisition against safety (Wilson et al., 2012). Canopy openings provide regenerating vegetation and patches of nutritionally valuable forage selected by browsing mammals (Kuijper et al., 2009; Wilson et al., 2024). The foraging value of beaver-created canopy openings was supported by our models, which showed that although ungulates visited beaver sites less frequently, they spent more time near water and were more likely to forage there. At the same time, beaver sites was associated with increased vigilance, while foraging probability declined with increasing large carnivore occurrence, suggesting that predation risk was indeed perceived by ungulates. Where resource availability and predation risk are spatially overlapped, successful foraging in suitable habitat patches comes along with modification in behaviour to avoid predators (Brown et al., 1999; Creel & Christianson, 2009; Schmidt & Kuijper, 2015). At beaver sites, this resource–risk trade-off was expressed not by avoidance of beaver-modified habitat, but spatially and temporally adjusted time budget to fit the fine-scale distribution of food resources and predation risk generated indirectly by beavers.

Although beaver presences is associated with large mammal space use and behaviour, the strength and direction of relationships with site type, distance from water, canopy openness and predator-prey interactions differed among seasons. Seasonal habitat selection by ungulates reflects temporal changes in the relative value of different habitats, as forage availability, cover, predation risk and energetic costs vary throughout the year (Owen-Smith et al., 2010). According to the risk-allocation hypothesis, antipredator behaviour should vary with the temporal context of risk and the ability of prey to afford its energetic costs (Lima & Bednekoff, 1999). Deer were more responsive to predation risk during milder parts of the year, whereas winter conditions reduced behavioural responsiveness as limited forage availability, poor energetic state and high movement costs constrained the ability to avoid predators (Clare et al., 2023). Similarly, in our study, greater large carnivore occurrence was associated with increased vigilance and reduced foraging in summer and autumn, but with reduced vigilance and increased foraging in winter and spring. Because beaver engineering is concentrated close to the shoreline, these modifications create relatively persistent and spatially predictable patches of resources and potential risk. However, the behavioural response of ungulates to increasing distance from the water changed through the year: for example, foraging was concentrated closer to water at beaver sites in summer and autumn, whereas in winter and spring vigilance was highest close to water and declined with increasing distance. Thus, the same beaver-modified part of the landscape may represent a profitable feeding patch in one season but a more constrained or risky environment in another. This spatially and temporally contingent response is consistent with the broader view of ecosystem engineering as a dynamic process rather than a fixed habitat effect (Sanders & Frago, 2024). The ecological effect of beaver engineering should therefore not be viewed as fixed, but as a dynamic contribution to ecosystem complexity, in which persistent habitat modification continually reshapes the spatial and temporal context of resource use, risk and species interactions.

## Conclusion

Our study demonstrates that the ecological consequences of Eurasian beaver recovery extend beyond habitat engineering. Through habitat-mediated pathways and changes in trophic interactions, beavers reorganize mammal communities not through species turnover, but through shifts in space use, behaviour and interactions among existing community. This broader perspective positions beaver not only as habitat modifier or facilitator of biodiversity, but also as integral participant of the ecological networks. Effects associated with beaver presence differed among trophic groups and emerged from the combined influence of habitat modification, direct associations with beavers and indirect relationships mediated by other mammals. In particular, beaver-modified habitats altered the ecosystem in which ungulates encountered both resources and risks. The strength and direction of beaver effects varied along the water-land gradient and among seasons, showing that the ecological role of beavers is not fixed. Beaver recovery may therefore restore more than habitat heterogeneity or individual trophic links, but also introduce spatial and temporal context in which assemblage of terrestrial mammals interact.

## CRediT authorship contribution statement

Izabela Fedyń: Project administration, Writing – original draft, Conceptualization, Methodology, Data curation, Investigation, Formal analysis, Visualization. Michał Ciach: Investigation, Writing – original draft, Supervision, Conceptualization.

## Declaration of Generative AI and AI-assisted technologies in the writing process

During the preparation of this work the authors used ChatGPT to proofread the text. After using this tool, the authors reviewed and edited the content as needed and take full responsibility for the content of the publication.

## Acknowledgements

We wish to express our gratitude to Piotr Kutrzeba, Wojciech Sobociński, Jakub Wyka and Karol Wróbel for their help with the fieldwork. This study was financially supported by the National Science Centre, Poland (grant No. 2021/41/N/NZ8/02423).

## Ethical statement

The study was performed in accordance with Polish law.

## Data availability

The data used in this study is available on request from the corresponding author.

## Supplementary materials

**Table S1.** Ethogram developed for behavioral classification of mammals pictures recorded by camera traps.

| Behaviour | Description |
| --- | --- |
| Feeding | The individual is observed actively consuming food or water, including grazing, browsing, foraging, chewing or drinking. The head is oriented toward the ground, vegetation, carcass, or water source, and movement is limited to feeding-related actions. |
| Movement | The individual is moving from one location to another. This includes walking, running, trotting, or bounding, with the body oriented in a consistent direction of travel. |
| Vigilance | The individual displays alert postures indicative of environmental monitoring, such as standing still with head raised, ears erect, scanning the surroundings, or frequently changing head orientation without locomotion. |
| Social behaviour | Interactions between two or more individuals, including interactions as play, hunting or mating behaviour, or territorial marking (e.g. scent marking, scratching). |
| Resting | The individual is lying or sitting, including sleeping or resting postures. |
| Unidentified behaviour | Behaviour could not be reliably classified due to incomplete visibility of the individual (e.g. only part of the body visible) or insufficient information to assign the observation to another category. |

**Table S2.** Number of photos, detections (time interval 30 min) and detections per 100 camera-days of ungulates, large carnivores, mesocarnivores, other wild mammals and other detections recorded by camera traps on sampling plots on Eurasian beaver (*Castor fiber*) sites (B; N = 450) and paired reference sites (R; N = 449) in Poland (central Europe).

| Group of mammals<br>Species |  | Number of<br>photos |  | Number of<br>detections |  | Detections per<br>100 camera-days |  |
| --- | --- | --- | --- | --- | --- | --- | --- |
|  |  | B | R | B | R | B | R |
| **Ungulates** |  |  |  |  |  |  |  |
| Moose | *Alces alces* | 106 | 174 | 33 | 51 | 0.45 | 0.70 |
| European bison | *Bison bonasus* | 571 | 284 | 13 | 29 | 0.18 | 0.40 |
| Roe deer | *Capreolus capreolus* | 5544 | 7014 | 970 | 1334 | 13.16 | 18.18 |
| Red deer | *Cervus elaphus* | 3570 | 4062 | 1075 | 1303 | 14.58 | 17.76 |
| Fallow deer | *Dama dama* | 15 | 77 | 6 | 17 | 0.08 | 0.23 |
| Wild boar | *Sus scrofa* | 612 | 737 | 124 | 158 | 1.68 | 2.15 |
| **Large carnivores** |  |  |  |  |  |  |  |
| Grey wolf | *Canis lupus* | 251 | 168 | 112 | 93 | 1.52 | 1.27 |
| Eurasian lynx | *Lynx lynx* | 14 | 8 | 8 | 5 | 0.11 | 0.07 |
| Brown bear | *Ursus arctos* | 131 | 9 | 30 | 5 | 0.41 | 0.07 |
| **Mesocarnivores** |  |  |  |  |  |  |  |
| European wildcat | *Felis silvestris* | 47 | 24 | 36 | 21 | 0.49 | 0.29 |
| Pine marten | *Martes martes* | 284 | 283 | 202 | 223 | 2.74 | 3.04 |
| European badger | *Meles meles* | 46 | 105 | 27 | 82 | 0.37 | 1.12 |
| Raccoon dog | *Nyctereutes procyonoides* | 31 | 23 | 20 | 18 | 0.27 | 0.25 |
| Red fox | *Vulpes vulpes* | 1627 | 539 | 421 | 413 | 5.71 | 5.63 |
| **Other wild mammals** |  |  |  |  |  |  |  |
| European beaver | *Castor fiber* | 1267 | 14 | 321 | 12 | 4.35 | 0.16 |
| European hare | *Lepus europaeus* | 22 | 59 | 17 | 44 | 0.23 | 0.60 |
| Eurasian otter | *Lutra lutra* | 13 | 15 | 8 | 7 | 0.11 | 0.10 |
| European polecat | *Mustela putorius* | 14 | 9 | 7 | 8 | 0.09 | 0.11 |
| Short-tailed weasel | *Mustela erminea* | 1 | 0 | 1 | 0 | 0.01 | 0.00 |
| Least weasel | *Mustela nivalis* | 0 | 3 | 0 | 3 | 0.00 | 0.04 |
| American mink | *Neovison vison* | 1 | 13 | 1 | 13 | 0.01 | 0.18 |
| Red squirrel | *Sciurus vulgaris* | 136 | 184 | 105 | 151 | 1.42 | 2.06 |
| Rodents |  | 38 | 54 | 33 | 37 | 0.45 | 0.50 |
| **Other detections** |  |  |  |  |  |  |  |
| Birds |  | 1094 | 405 | 531 | 271 | 7.20 | 3.69 |
| Domestic dog | *Canis familiaris* | 129 | 91 | 34 | 42 | 0.46 | 0.57 |
| Domestic cat | *Felis catus* | 3 | 14 | 2 | 8 | 0.03 | 0.11 |
| Human |  | 1440 | 1347 | 315 | 353 | 4.27 | 4.81 |
| Vehicle |  | 43 | 51 | 13 | 12 | 0.18 | 0.16 |

**Table S3.** Means of the habitat variables measured on sampling plots on Eurasian beaver (*Castor fiber*) sites (N = 450) and paired reference sites (N = 449) in Poland (central Europe). Differences tested with Student t-test.

| Variable | mean Beaver sites | mean Reference sites | difference between Reference-Beaver sites (%) | p |
| --- | --- | --- | --- | --- |
| Canopy closure | 46.3 | 59.6 | -22.3 | 0.000 |
| Coverage of undergrowth | 22.3 | 21.9 | 1.8 | 0.771 |
| Number of snags | 3.6 | 2.3 | 56.5 | 0.005 |
| Number of logs | 8.1 | 4.2 | 92.9 | 0.000 |
| Number of uproots | 0.3 | 0.4 | -25.0 | 0.148 |
| Forest floor diversity | 1.1 | 1.0 | 10.0 | 0.041 |
| Coverage of grass | 26.6 | 14.0 | 90.0 | 0.000 |
| Coverage of herbs | 20.4 | 24.0 | -15.0 | 0.025 |
| Coverage of blueberry | 5.3 | 3.5 | 51.4 | 0.043 |
| Coverage of blackberry | 10.9 | 11.9 | -8.4 | 0.466 |

**Fig. S1.**
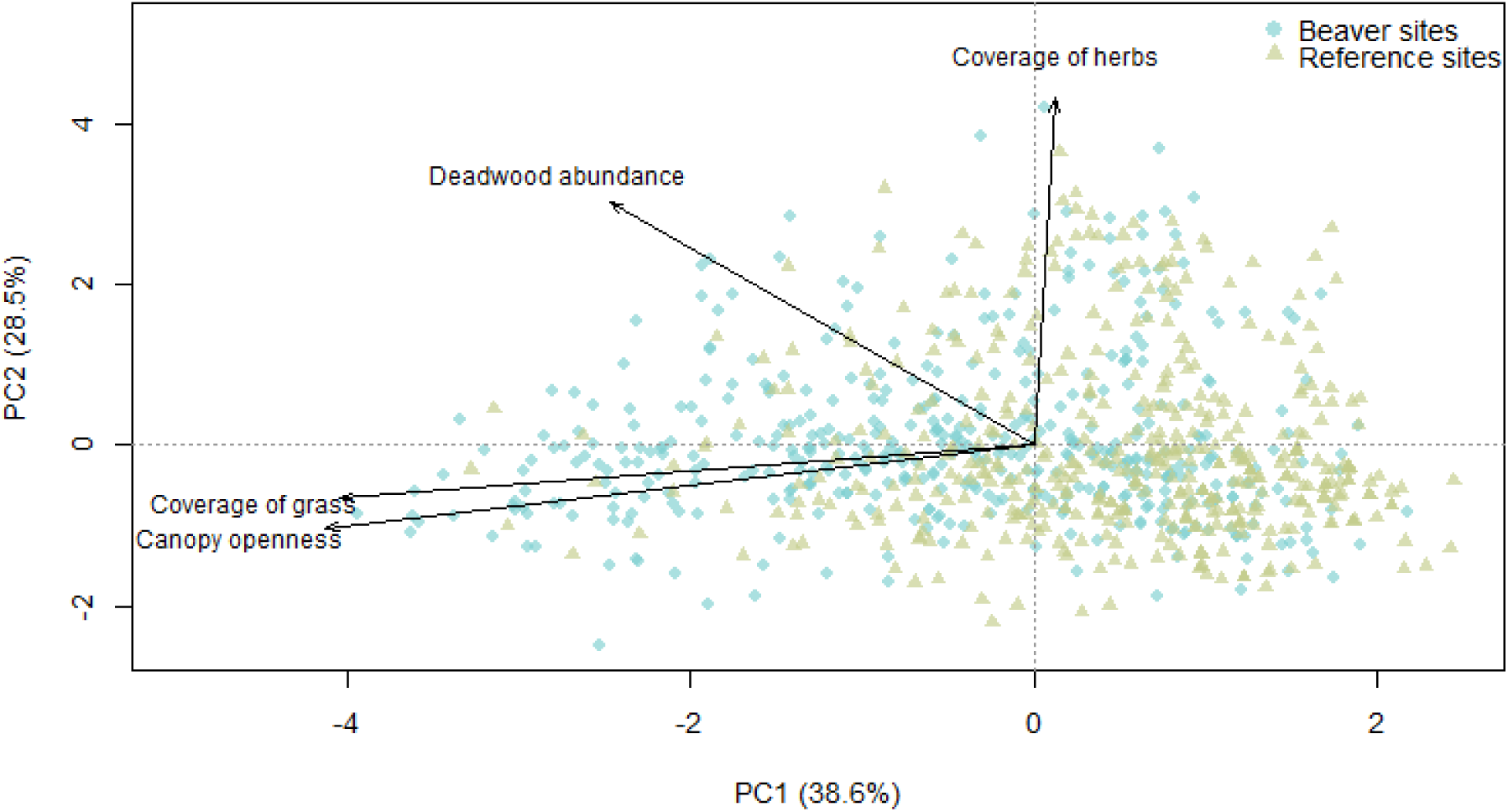
Principal component analysis build based on habitat variables on sample plots on Eurasian beaver (*Castor fiber*) (N = 450) and reference (N = 449) sites in Poland (central Europe).

**Table S4.** Mixed-effects model results for the effects of site type, distance to water and their interaction on canopy openness and deadwood availability at Eurasian beaver (*Castor fiber*) sites and paired reference sites in Poland (central Europe).

|  | Variable | Estimate | SE | df | t | p |
| --- | --- | --- | --- | --- | --- | --- |
| Canopy openness | Intercept | -0.27 | 0.11 | 58.04 | -2.57 | 0.013 |
|  | Type | 0.52 | 0.15 | 57.89 | 3.49 | 0.001 |
|  | Distance | -0.04 | 0.04 | 821.50 | -1.10 | 0.270 |
|  | Type:Distance | -0.35 | 0.05 | 821.02 | -6.68 | 0.000 |
| Deadwood | Intercept | -0.30 | 0.12 | 57.73 | -2.56 | 0.013 |
|  | Type | 0.61 | 0.17 | 57.63 | 3.66 | 0.001 |
|  | Distance | -0.11 | 0.03 | 817.00 | -3.22 | 0.001 |
|  | Type:Distance | -0.30 | 0.05 | 816.63 | -6.15 | 0.000 |

**Table S5.** Mixed-effects model results for the effects of site type, distance to water and their interaction on number of detections of large carnivores, mesocarnivores and ungulates at Eurasian beaver (*Castor fiber*) sites and paired reference sites in Poland (central Europe).

| Group of mammals | Variable | Estimate | SE | z | p |
| --- | --- | --- | --- | --- | --- |
| Large carnivores detections | Intercept | -5.31 | 0.25 | -21.61 | 0.000 |
|  | Type | 0.49 | 0.26 | 1.85 | 0.065 |
|  | Distance | -0.01 | 0.13 | -0.04 | 0.964 |
|  | Type:Distance | -0.11 | 0.17 | -0.64 | 0.522 |
| Ungulates detections | Intercept | -1.11 | 0.06 | -17.50 | 0.000 |
|  | Type | -0.29 | 0.09 | -3.30 | 0.001 |
|  | Distance | 0.08 | 0.04 | 2.06 | 0.040 |
|  | Type:Distance | 0.24 | 0.06 | 4.26 | 0.000 |
| Mesocarnivores detections | Intercept | -2.39 | 0.10 | -24.61 | 0.000 |
|  | Type | -0.16 | 0.13 | -1.27 | 0.203 |
|  | Distance | -0.28 | 0.08 | -3.49 | 0.000 |
|  | Type:Distance | 0.29 | 0.11 | 2.65 | 0.008 |

